# Relationships between landscape structure and the prevalence of two tick-borne infectious agents, *Anaplasma phagocytophilum* and *Borrelia burgdorferi* sensu lato, in small mammal communities

**DOI:** 10.1101/718403

**Authors:** Grégoire Perez, Suzanne Bastian, Amélie Chastagner, Albert Agoulon, Yann Rantier, Gwenaël Vourc’h, Olivier Plantard, Alain Butet

**Author notes:** UMR BIPAR, INRA-Anses-ENVA, 94701 Maisons-Alfort, France. ANSES, Laboratoire de Ploufragan-Plouzané-Niort, 22440 Ploufragan, France (Amélie Chastagner).

## Abstract

**Context:** By modifying ecosystems, land cover changes influence the emergence, the spread and the incidence of vector-borne diseases.

**Objective:** This study aimed at identifying associations between landscape structure and the prevalence of two tick-borne infectious agents, *Anaplasma phagocytophilum* and *Borrelia burgdorferi* s.l., in small mammal communities.

**Methods:** Small mammals were sampled in 24 sites along a gradient of woodland fragmentation and hedgerow network density, and screened for infectious agents with real-time PCR techniques. For each site, structural variables (composition and configuration) of the surrounding landscape at various scales (0-500 m) and variables of wooded habitats connectivity based on graph theory and least cost path distances for the two dominant species, bank voles (*Myodes glareolus*) and wood mice (*Apodemus sylvaticus*), were computed.

**Results:** The *A*. *phagocytophilum* prevalence increased with wooded habitats cover (0-500 m), likely through host population size, and increased slightly with bank vole abundance, which has a higher reservoir competence than wood mouse. The *B. burgdorferi* s.l. prevalence increased with wooded ecotones only at local scales (50-100 m). Wooded habitats connectivity measures did not improve models built with simple land cover variables. A more marked spatial pattern was observed for the prevalence of *A*. *phagocytophilum* than *B. burgdorferi* s.l..

**Conclusions:** This study highlights the interest of considering together the ecology of infectious agents (e.g. host specificity) and the host species community ecology to better understand the influence of the landscape structure on the spatial distribution of vector-borne infectious agents.

## Introduction

Landscape changes are suspected to cause infectious disease emergence or reemergence worldwide (Jones et al. 2008). Several authors depicted the complexity of the interactions between the landscape, hosts, vectors and humans, which drive the emergence of these diseases (Lambin et al. 2010). The spread of infectious agents in ecosystems relies partly on their life history traits shaped by their evolutionary histories (e.g. transmission modalities). Because species within communities exhibit various levels of reservoir or vector competence, the transmission depends on hosts and vectors diversity and abundance (Keesing et al. 2009).

Landscape structure (composition and configuration), by filtering and regulating richness, abundance and dispersal of species in hosts and vectors communities, may influence the transmission rate and the diffusion of infectious agents, which ultimately determine their distribution and prevalence (Suzán et al. 2015). For instance, an increase in the proportion of non-reservoir hosts may induce an increased number of missed transmissions between reservoir hosts, or an increased number of wasted bites (failed transmission events) by vectors. This may reduce the number of transmission events and has a so called “dilution effect” on prevalence (Ostfeld and Keesing 2000; Clay et al. 2009). Conversely, an increase in the proportion of competent hosts may increase the number of transmission events or efficient bites by vectors (Roche et al. 2013). When less competent species are more prone to local extinction, this non-random species loss in host communities seems to be a driver of the increased direct and vector-borne infectious disease transmission (Ostfeld and LoGiudice 2003; Johnson et al. 2013; Roche et al. 2013).

Habitat connectivity partly shapes hosts community structure. Large-bodied species, with small clutch/litter size and long gestation/incubation time are supposed to be more sensitive to habitat fragmentation because they experience higher local extinction risk. Conversely, small-bodied species, with large clutch/litter size and short incubation/gestation time can persist in fragmented habitat landscapes. These species need less habitat surface and benefit from a competition and predation release in these landscapes that are less favorable to predators and specialist species (Nupp and Swihart 1998, 2000; Gehring and Swihart 2003; Ferrante et al. 2017). As it happens, the latter kind of species also seem to be better reservoirs for infectious agents (Gottdenker et al. 2012; Huang et al. 2013; Ostfeld et al. 2014). Thus, in fragmented habitat landscapes, one could expect an amplification of infection prevalence supported by a higher density of more competent hosts (Allan et al. 2003; Rubio et al. 2014). However, habitat connectivity can also indirectly influence the risk of local extinction of infectious agents by shaping hosts – and vectors – population size, as demonstrated on Hantavirus in bank voles (Guivier et al. 2011). Land use is highly changing in Europe (agricultural intensification in some areas and rewilding in others; Reidsma et al. 2006). Thus, more studies explicitly assessing how landscape composition, configuration, and the resulting functional connectivity influence interactions and abundances of hosts, vectors and infectious agents to better understand infectious diseases risk are needed.

Small mammals are reservoir hosts of many tick-borne infectious agent taxa: protozoan (e.g. *Babesia* sp., Hersh et al. 2012), bacteria (e.g. *Borrelia* sp., Gern et al. 1998; *Anaplasma* sp., Stuen et al. 2013) and viruses (e.g. tick-borne encephalitis virus, Mansfield et al. 2009). Small mammal communities composition is known to vary according to landscape features like habitat patch size, shape and isolation (Nupp and Swihart 2000; Michel et al. 2007) and to the constraints that the landscape matrix imposes on the displacements of small mammals (Szacki et al. 1993).

Tick displacements *per se* are reduced and their dispersal occurs mostly while feeding on their hosts (Araya-Anchetta et al. 2015). Hard ticks (Arthropoda, Arachnida, Ixodidae) need to have a first blood meal on a vertebrate host to molt from larva to nymph, a second one to molt from nymph to adult then adult females need a third one to produce eggs once fertilized by adult males. Tick-borne infectious agents can be acquired by ticks or transmitted to hosts at each blood meal. They disperse either by the dissemination of ticks getting infected while feeding on infected hosts, or by already infected ticks while attached on hosts. The dispersal distance of infectious agents depends on home ranges and competences of tick hosts (Kurtenbach et al. 2002b). Thus, landscape features affecting host communities and/or their dispersal are of paramount importance in the distribution and the local prevalence of tick-borne infections.

This study focused on *Anaplasma phagocytophilum* and *Borrelia burgdorferi* sensu lato (s.l.). The *A. phagocytophilum* bacteria ecotype associated to small mammals is transmitted mostly by *I. trianguliceps* and maybe other endophilic (burrow dwelling) tick species (Bown et al. 2009; Blaňarová et al. 2014; Jahfari et al. 2014). Infections in small mammals last about a couple of months (Bown et al. 2003). No transovarial transmission (from engorged females to their offspring) of these bacteria is known in *Ixodes* ticks (Stuen et al. 2013).

In Europe, several genospecies of the *B*. *burgdorferi* s.l. complex can be hosted by small mammals: *B. afzelli, B. bavariensis* (formerly *garinii* OspA serotype 4 strain), *B*. *bissetti, B. burgdorferi* sensu stricto (s.s.), and *B*. *spielmani* (Kurtenbach et al. 2002a, 2006; Margos et al. 2009; Coipan and Sprong 2016). However, host ranges of some of these genospecies include also larger mammals like hedgehogs *(Erinaceus europaeus)* and squirrels *(Sciurus vulgaris)* (Skuballa et al. 2012; Pisanu et al. 2014). The *B*. *burgdorferi* s.l. bacteria can be transmitted by several tick species, but the exophilic (questing for host on the vegetation) tick species *I*. *ricinus* is assumed to be its main vector in Europe (Rizzoli et al. 2011). Small mammals are major hosts for *I*. *ricinus* larvae and only occasional hosts for nymphs, but they also host endophilic tick species like *I*. *trianguliceps* and *I*. *acuminatus* at all life stages (Boyard et al. 2008; Bown et al. 2008; Hofmeester et al. 2016; Perez et al. 2017). Therefore, the maintenance of the zoonotic cycle of these bacteria may partly rely on other tick species (Hubbard et al. 1998; Heylen et al. 2013; Szekeres et al. 2015). Infections in small mammals are lifelong (Gern et al. 1994; Humair et al. 1999). Transovarial transmission is assumed to be null or rare (Rollend et al. 2013).

Here, we investigated at a local scale how landscape composition, configuration, and resulting habitat connectivity for small mammals, directly or *via* relationships with species abundances, can explain the prevalence of *A*. *phagocytophilum* and *B*. *burgdorferi* s.l. in small mammals. We specifically tested the three following hypothesis on the prevalence drivers:

(**H1**) Host population size: prevalence would be higher in sites supporting larger host populations (i.e. in wooded habitat patches either of large area, with more surrounding wooded habitats and/or more connected).
(**H2**) Overall host community competence: the landscape would indirectly influence the prevalence by acting on the relative abundance of host species exhibiting different reservoir competence, modifying the overall community competence.
(**H3**) Infectious agents’ specificity: a stronger relationship between the prevalence and landscape variables would be observed for the *A*. *phagocytophilum* ecotype specific to small mammals, than for the *B*. *burgdorferi* s.l. group, which shows a wider range of host and of vector species.

## Materials and Methods

### Study area

The study took place in the ‘Zone Atelier Armorique’ (Brittany, France), a 132 km^2^ Long-Term Ecological Research (LTER) area labeled by the CNRS (Centre National de la Recherche Scientifique) and belonging to the International LTER network. This area includes different landscapes with various land uses, from deciduous mixed forest, to livestock-crop mixed farming landscapes, to cereal-oriented (mainly cereals and maize) open farming landscapes. Compared to mixed farming landscapes, open farming landscapes have fewer (and smaller) woodlots (area: 10% vs 15%), fewer grasslands (area: 20% vs. 40%), larger crops patches (mean area: 2.3 ha vs. 1.3 ha), and looser hedgerow networks (50 vs. 115 m/ha) (Millán de la Peña et al. 2003; Al Hassan et al. 2012; Perez et al. 2016).

### Sampling strategy

Wooded habitats support higher abundances of small mammals, better abiotic conditions for ticks (moisture) and are as such supposed to be key habitats for small mammals-ticks interactions (Boyard et al. 2008). For these reasons, 24 sampling sites were selected in wooded habitat patches in various landscape contexts (Perez et al. 2016). The 24 sampling sites, at least 500 m apart from each other to ensure their spatial independence, were placed as follows: six in the forest core; six in forest edge landscapes; six in mixed farming landscapes; and six in open farming landscapes. In each of the two agricultural landscape types, three sites were selected along hedgerows (six sites overall) and three along woodlots edges (six sites overall). At forest edges and in agricultural landscapes, the sampling sites (n = 18) were selected at the ecotones between wooded habitats and grasslands, as meadows are less unfavorable to ticks than cultivated plots. Because small mammals start breeding in spring and populations peak generally in autumn, we sampled in May-June and in October in 2012 and 2013. These periods of the year also correspond to the main and the secondary activity peak of *I. ricinus* nymphs, respectively, and conversely for *I*. *trianguliceps* adult females, while the nymphs of this latter species are active from summer to mid-autumn (Randolph 1975a).

### Small mammal sampling and ethic statement

Small mammals were trapped using French model (INRA) live traps with wooden dormitory boxes. These traps can catch small mammals from 4 to 40 grams, including shrews and small rodents. Trap-lines (100 m-long) were constituted of 34 traps (three meters apart) baited with a mix of commercial seeds for pet rodents (sunflower, wheat, etc.), dry cat food, and a piece of apple for water supply. Traps were checked in the morning after 24 and 48 hours. Animals were brought to the field lab to be identified at the species level, euthanized by pentobarbital injection, weighted, sexed and dissected.

Traps were designed to limit as much as possible the stress or injury of the animals. The bait was designed to obtain an optimal catch rate across small mammal species and a good survival of individuals. Targeted animals were not protected species and thus no special authorization was needed, according to the French law in force. All individuals were euthanized by authorized experimenters according to current French law and to the European guidelines on the use of animals in science (Close et al. 1997).

### Molecular detection of infectious agents

DNA was extracted from small mammal spleens and ears with Macherey-Nagel NucleoSpin Tissue kits (Chastagner et al. 2016). The screening for *A. phagocytophilum* was performed on DNA extracts from small mammal spleens by real-time PCR targeting the *msp2* genes according to the protocol of Courtney et al. (2004). The screening for *B*. *burgdorferi* s.l. was performed on DNA extracts from small mammal ears by real-time PCR in SybrGreen according to the protocol of Halos et al. (Halos et al. 2010). The *B*. *burgdorferi* s.l. prevalence was low and, for technical reasons, identifying them to the geno-species level (genetically discriminated *Borrelia* species) was not possible (data not shown). Thus, all the *B*. *burgdorferi* s.l. genospecies were combined hereafter. Although all geno-species which can infect small mammals do not share the same host range, *B*. *afzelii* is known to be predominant in the captured species (Kurtenbach et al. 2002a; Marsot et al. 2013). Therefore, this simplification is assumed not to bias results. As temporal the variation of prevalence has been studied elsewhere (Perez et al. 2017), the data from all the sampling sessions were grouped for each site to study only the spatial variation hereafter.

### Landscape variables

#### Landscape data

The used landscape data were: a shapefile of the land cover of 2012 kindly provided by the ‘LETG-COSTEL-Rennes’ lab *via* the ‘Zone Atelier Armorique’; a road network shapefile provided by the ‘Institut National de l’Information Géographique et Forestière’ (IGN); and a 5 m-resolution raster file of the land cover of 2010 from Gil Tena et al. (2014) for woodland and hedgerows. The land covers were characterized as “woodland”, “hedgerows” (abandoned lands and hedgerows extracted from the 5 m-resolution raster file), “grassland”, “crops”, “roads” (ranked according to the IGN’s ‘road importance index’), “buidings” and “water areas”. All files were aggregated with ArcGIS 10.3.1 for Desktop, Esri ®, into a single 5 m-resolution raster file that was used in subsequent analyses.

#### Landscape composition and configuration variables

The proportion of wooded habitats (‘Wooded’ hereafter), computed as the proportion of “woodland”/“hedgerows” pixels, was chosen as a landscape composition variable representative of the amount of favorable permanent habitats surrounding trap-lines. The proportion of wooded ecotones surrounding trap-lines (‘Ecotones’ hereafter), computed as the proportion of ‘wooded habitat’ and “grassland”/”crops”/“roads” adjacent pixels (pixel couples), was chosen as a landscape configuration variable. Such ecotones are known to be favorable to generalist species and are places where species from different habitat can co-occur (Meunier et al. 1999; Ouin et al. 2000; Boyard et al. 2008). This configuration variable depends on landscape heterogeneity, and is maximal in landscapes with moderate woodland fragmentation.

The relationships between landscape variables and small mammal abundances or infection prevalence were evaluated at various spatial scales by computing these variables in circular zones of various radii around trap-line centres with Chloe2012 software (Boussard and Baudry 2014). Because trap-lines were 100 m-long, the first circular 50 m-radius zones were considered as informative on trapping habitats only (‘local’). Larger circular zones with an additional 50, 100, 200, and 500 m radius were considered.

#### Landscape habitat connectivity metrics

The dispersal of the studied infectious agents is known to rely mostly on larval ticks, which acquire them by feeding on an infected small mammal and falling off in the latter’s home range. The dispersal of infectious agents by young small mammals, which have not yet been highly exposed to ticks and are thus yet unlikely infected, can be considered as negligible (Randolph 1975b; Sinski et al. 2006; Kallio et al. 2014; Perez et al. 2017). Furthermore, despite *I*. *ricinus* nymphs can disperse bacteria on longer distances attached on hosts with larger home ranges, small mammals are unlikely to be infected by *I*. *ricinus* adult females because they rarely host them. For these reasons, we performed the functional connectivity analysis at the scale of small mammals’ home ranges.

Thus, we modelled the functional connectivity of the landscape for small mammals to evaluate its relationship with their infection prevalence. We used the difference in Probability of Connectivity metric (‘dPC’), which is an integrative measure of reachable habitats based on a negative exponential function (Saura and Pascual-Hortal 2007). The three dPC fractions were also used separately: the intra-patch connectivity (‘dPCintra’; no inter-patch displacement/dispersal), which is dependent of patch area only, so this latter variable was used instead for simplification purpose (‘Area’); the inter-patch connectivity (‘dPCflux’) and the “stepping stone” connectivity (‘dPCconnector’; connecting other patches).

Land cover cost files were built only for the two dominant rodent species with contrasted habitat preference: wood mice and bank voles (Boyard et al. 2008). Because these species move preferentially along linear structures or across the shortest path between suitable habitat patches (Zhang and Usher 1991), the functional distance between two habitat patches was set using least cost path distances. For “woodland”, “grassland” and “crops”, costs were assigned like the inverse of the relative species abundance from the literature (Ouin et al. 2000; Tattersall et al. 2002; Boyard et al. 2008; Vourc’h et al. 2008). The reference cost for “woodland” was one. For wood mice and bank voles, the chosen costs were respectively four and fifteen for “grassland”, and five and fifteen for “crops”. A maximum cost of 100 was assigned arbitrary to “water areas” and “buildings”. Roads are repulsive habitats for small mammals because they impede their displacements (Mader 1984; Rico et al. 2007). They also cause a traffic-related mortality (Ruiz-Capillas et al. 2015). Eskilkdsen (2010) reported that finding bank voles were 15 times less likely on the opposite side of a barrier (forest path or road) than on the same side. Thus, a gradient of costs from 15 (countryside and forest tracks) to 75 (expressways) was assigned to “roads” according to the ‘road importance index’.

The median distance travelled by an individual was modelled using negative exponential functions based on data from the same landscapes (Papillon et al. 2002), and from a forest landscape of the Berkshire, United-Kingdom (Kikkawa 1964). In consistency with the lower mobility of bank voles compared to wood mice, these distances were estimated at 25 and 100 m, respectively (Zhang and Usher 1991). The nodes were defined as patches with an area of at least 0.025 ha, corresponding to the minimum home range size of the studied species (Kikkawa 1964). Graphs were created based on least cost path networks for each species. To compute connectivity measures with ‘isolation by distance’ models, graphs based on Euclidian distance were also created for displacement/dispersal distances corresponding to half the radius used to compute land cover variables: 25, 50, 100, and 250 m. Nodes were adjacent woodland habitats pixels (“woodland” and “hedgerows”).

Connectivity measures were computed at the patch level for each graph. Because habitat patch areas and ‘dPC’ values were over-dispersed, those variables were log-transformed hereafter. Least cost paths, graphs, and connectivity measures were computed using Graphab 1.2.3 (Foltête et al. 2012). An example of resulting graphs is shown in **Appendix 1**. A summary of all the landscape variables is shown in **Table 1**.

**Appendix 1:**
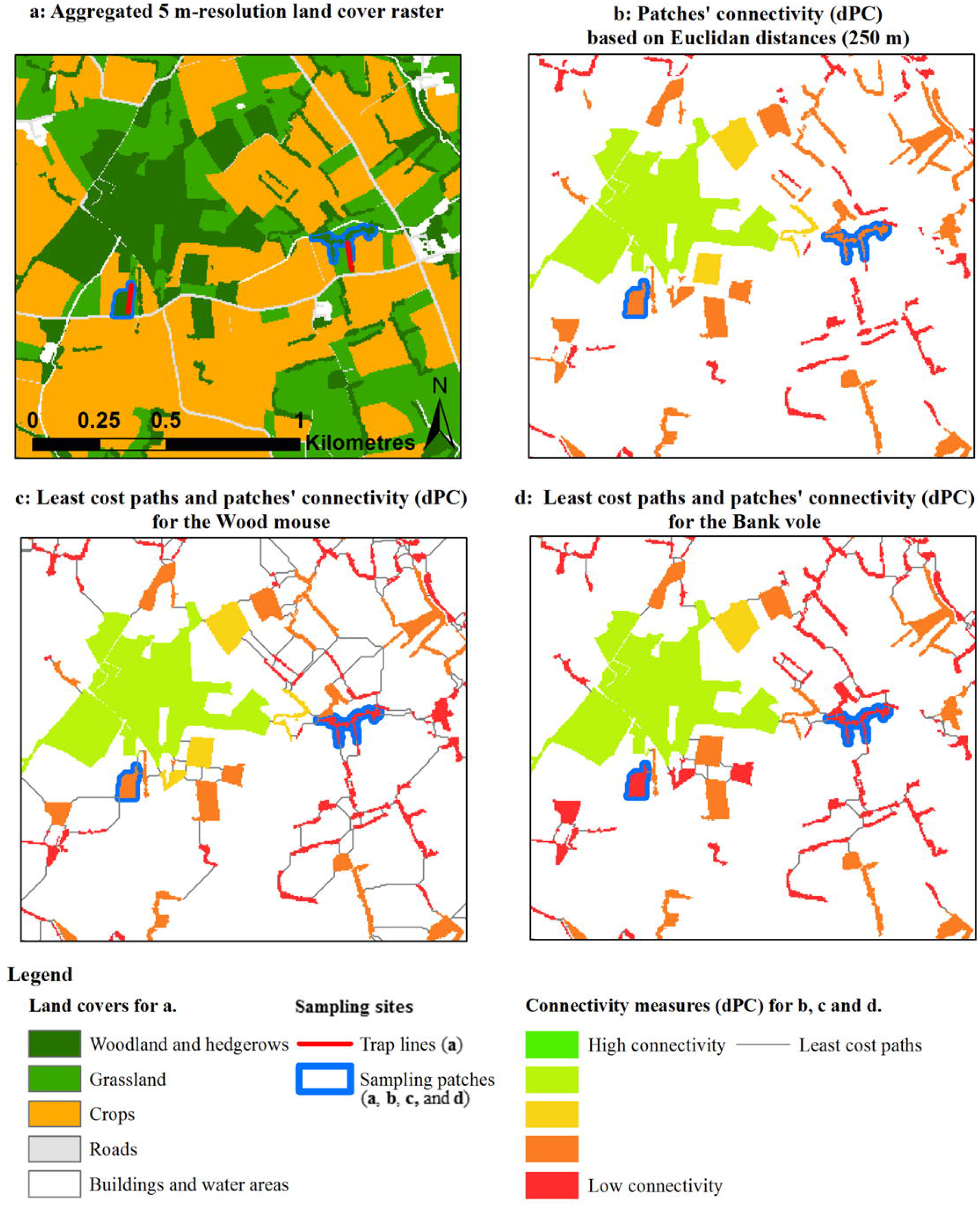
Example of the habitats landscape connectivity analyses. Final aggregated land cover 5 m-resolution raster file (**a**) and ‘dPC’ (‘difference in Probability of Connectivity’) metric computed on different graphs (**b**, **c**, and **d**). The sampling patch on the right (surrounded in blue) appears moderately connected with Euclidian distances (**b**), while it is weakly connected when weighted by least cost paths (**c** and **d**) because it is surrounded by a river at North and by roads on other sides (see **a**). The sampling patch on the left is moderately connected for the wood mouse (**c**) while it is weakly connected for the bank vole (**d**) because it is separated to other wooded habitat patches by large grassland or crops patches resulting in fewer connections for this latter species. See **Materials and Methods** for more details.

**Table 1:**
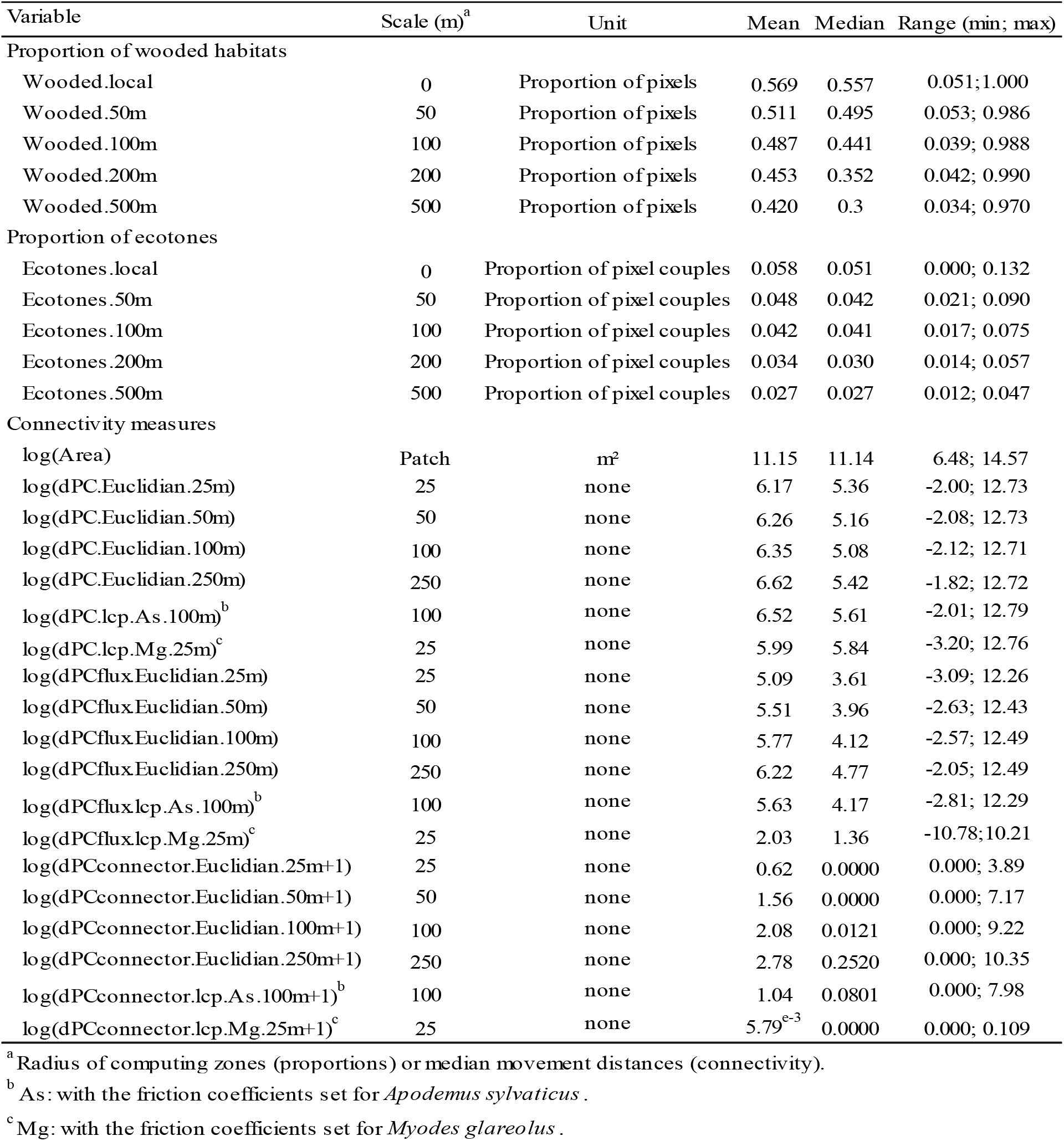
Landscape variables used in the study for the model selection procedures.

### Data analyses

To determine whether prevalence in wood mice and bank voles should be modelled separately, we first tested, for each infection, the correlation between prevalence in each species and the correlation between numbers of infected individuals of each species. Because of the non-normal distribution of prevalence data (Shapiro-Wilk test: p < 0.05), we used Spearman’s correlation tests.

To assess indirect effects of landscape structure on prevalence *via* the abundances of small mammal species, the relationship between these latter and landscape variables was tested. First, to determine whether wood mouse and bank vole abundances could be modelled separately, their correlation was tested through Pearson’s correlation test. Then, to choose the most appropriate error distribution of species abundances, Akaike Information Criterion corrected for finite sample size (AICc; Hurvich and Tsai 1989) values of the null models fitted with a normal error distribution and with a Poisson error distribution were compared. The normal error distribution had a better support (lower AICc-value) for wood mouse abundance (AICc-normal = 180.2, AICc-Poisson = 232.0) and for bank vole abundance (AICc-normal = 141.6, AICc-Poisson = 154.9). Species abundances were thus modelled using Linear Models (LMs).

The variables were first selected in single explanatory variable models (p < 0.1). Then, multiple explanatory variable LMs were built with the selected variables. To avoid collinearity problems, models with correlated variables (|r| > 0.5) were excluded. The models were then ranked based on AICc values, and the significance of the variable in the best models (ΔAICc < 2) was evaluated with type II ANOVAs (p < 0.05).

To assess whether prevalence were related to landscapes variables or species abundances, prevalence in small mammals per site was modelled using binomial Generalized Linear Models (GLMs) with the same explanatory landscape variables as above, with addition of wood mouse abundance (N.As), bank vole abundance (N.Mg), and the ratio between the bank vole abundance and the wood mouse abundance (ratio.Mg:As). Abundances were expressed as the total number of captured individuals per site. The variables were first selected in single explanatory variable models (p < 0.1). Another AICc-based model ranking and variable significance evaluation was finally performed on multiple explanatory variable GLMs, as described above.

All statistical analyses were performed using R software Version 1.2.5001 (R Development Core Team 2019). The R-package ‘car’ was used for ANOVAs, ‘MuMIn’ for model selections, and ‘psych’ for pairwise correlation tests between explanatory variables. The absence of autocorrelation in the dependent variables was checked using the ‘correlog’ function of the ‘ncf’ R-package.

## Results

A total of 612 small mammals belonging to five species were caught during the two sampling years (see **Table 2** for detailed results; the whole data set is available in supplementary material). Wood mice *(Apodemus sylvaticus)* and bank voles *(Myodes glareolus)* were largely dominant (74.2% and 24.3% of captured animals, respectively) and found in all the 24 sites. We captured nine individuals of three other species on seven sites: four field voles (*Microtus agrestis*) on three sites, two common pine voles (*Microtus subterraneus*) on two sites, and three crowned shrews (*Sorex coronatus*) on three sites. A maximum of four species were present in a single forest core site with all species excepting the common pine vole.

**Table 2:**
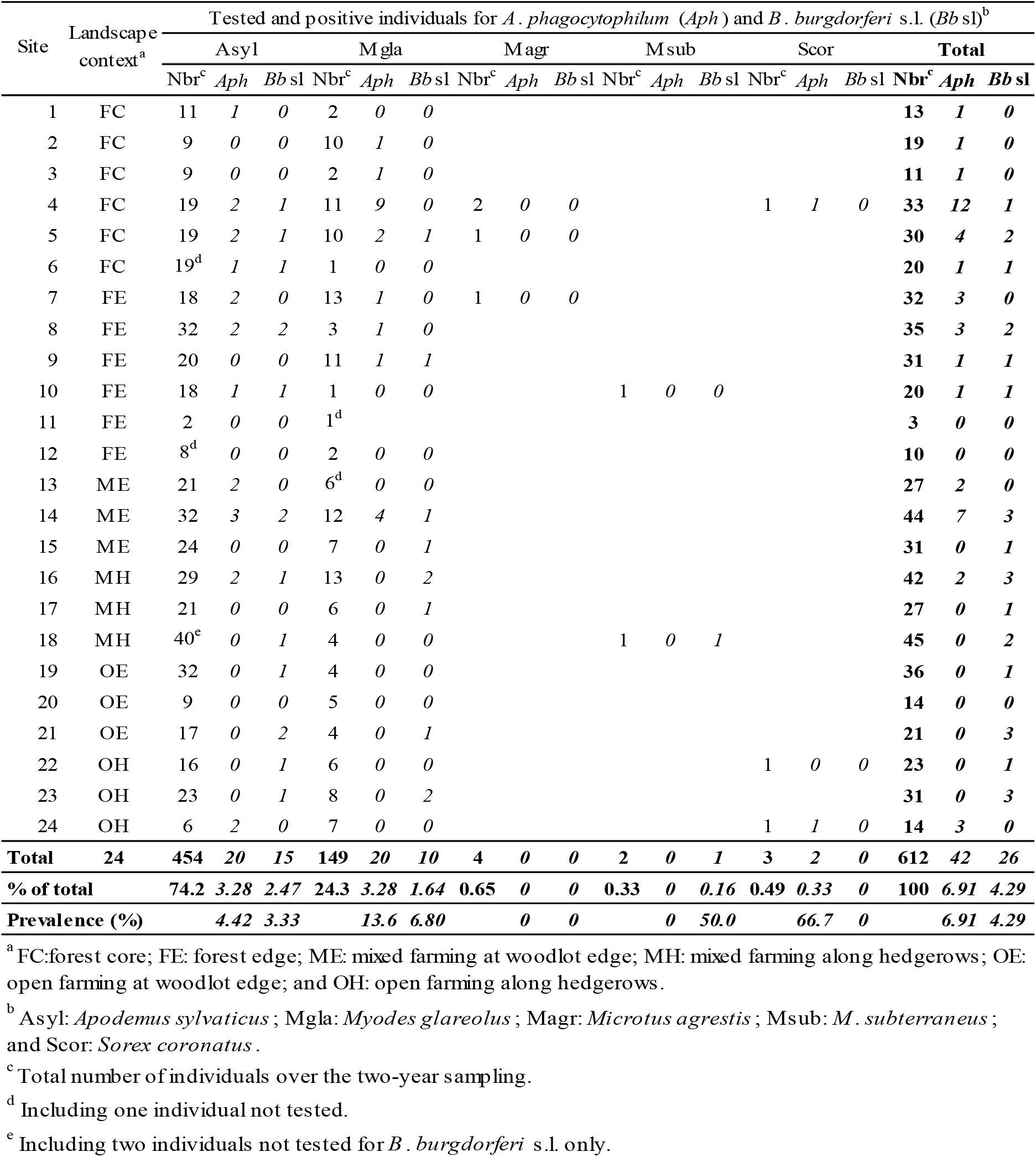
Results of *Anaplasma phagocytophilum* and *Borrelia burgdorferi* s.l. PCR detection per site over the two-year sampling (2012-2013).

Wood mouse abundance and bank vole abundance were not correlated (rho = 0.229, p = 0.281) and thus have been analysed separately hereafter. The best model for wood mouse abundance included only the proportion of ecotones in 500 m-radius zones (R2 = 0.290, p = 0.003) with a positive relationship. Bank vole abundance was not significantly related to any landscape variable (p > 0.05).

### *Results for* A. phagocytophilum

The analyses of *A*. *phagocytophilum* prevalence were based on the PCR results for 452 wood mice, 147 bank voles, four field voles, two common pine voles and three crowned shrews. Twenty wood mice, twenty bank voles and two crowned shrews were positive (**Table 2**). The number of positive wood mice and bank voles per site were significantly correlated (rho = 0.425, p = 0.038). However, the prevalence were not (rho = 0.291, p = 0.177). The results of the selection procedure of the GLMs of *A. phagocytophilum* prevalence as a function of landscape variables and species abundance variables are detailed in **Table 3**.

**Table 3:**
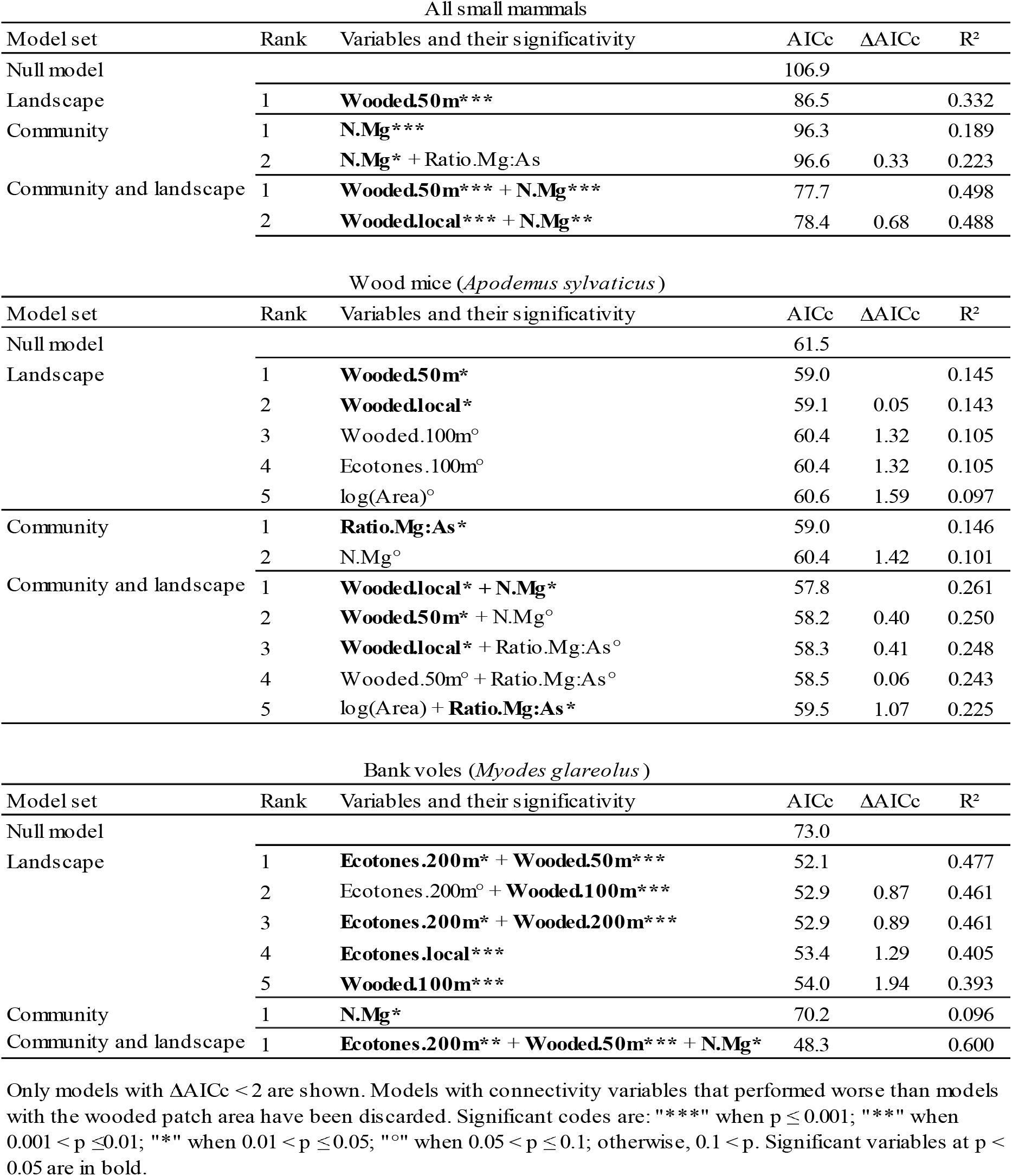
Best binomial Generalized Linear Models of *Anaplasma phagocytophilum* prevalence as a function of landscape variables and species abundance variables according to the AICc-based selection procedure.

The most supported GLM of *A*. *phagocytophilum* prevalence in all small mammals included bank vole abundance, (**Figure 1a**), and the proportion of wooded habitats in 50 m-radius zones, both displaying a significant positive relationship with the prevalence (**Figure 1a** and **1b**, respectively). The most supported GLM of prevalence in wood mice included bank vole abundance and the proportion of wooded habitats in the trapping habitats, both displaying a significant positive relationship (**Figure 1c** and **1d**). The ratio between bank vole and wood mouse abundances was significantly positively related to the prevalence in wood mice in the univariate model (not shown). The most supported GLM of prevalence in bank voles included the bank vole abundance, the proportion of wooded habitats in 50 m-radius zones, and the proportion of ecotones in 200 m-radius zones, all displaying a significant positive relationships with the prevalence (**Figure 1e**, **1f**, and **1g**, respectively).The relationship between prevalence and ecotones was negative in single explanatory variable models considering all sites, but positive when excluding forest sites (not shown). Thus, the apparent contradiction between single and multiple explanatory variables models likely results from the reduction of ecotones length in highly wooded landscapes (see the prevalence as a function of both landscape variables in **Figure 1h**).

**Figure 1:**
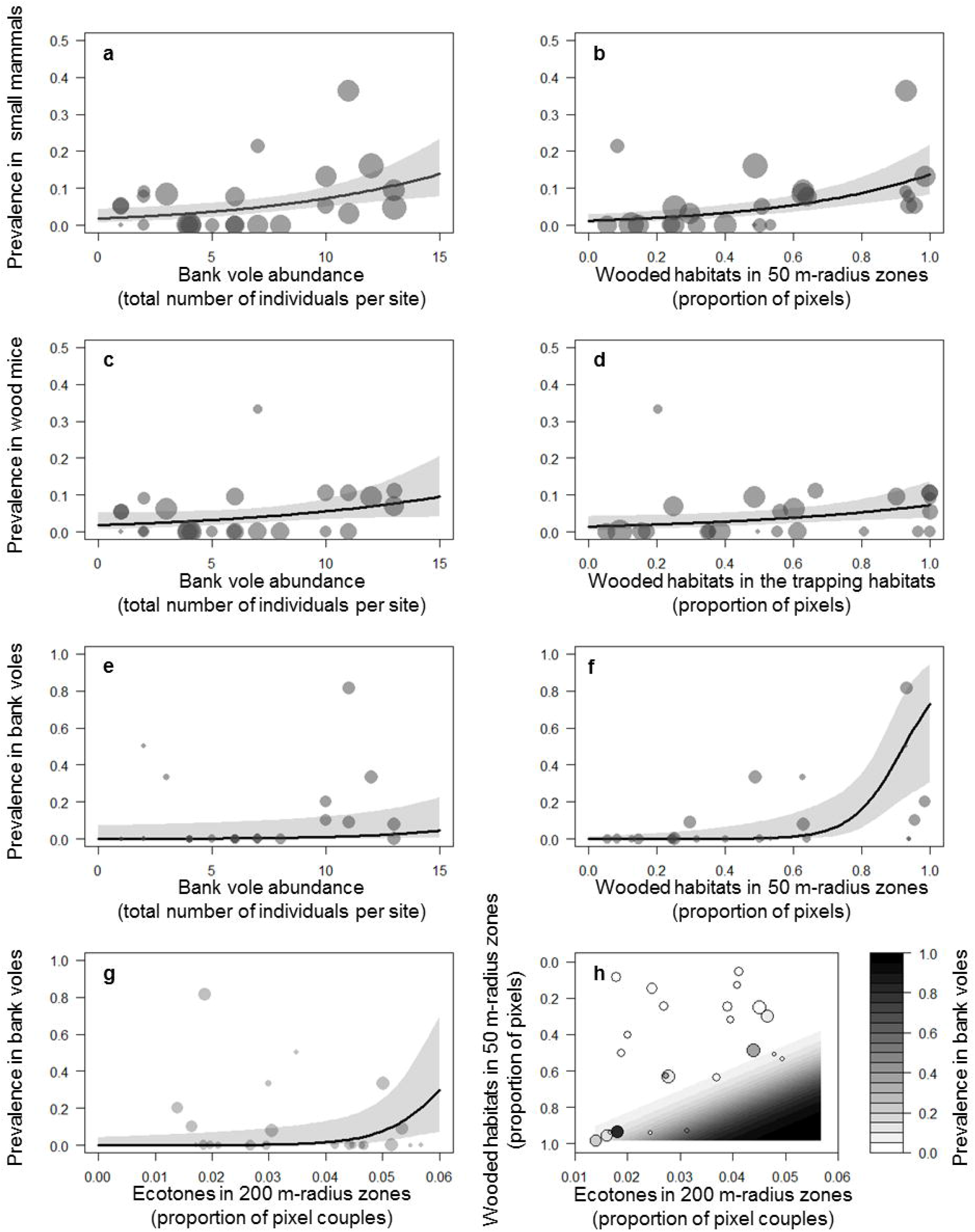
The *Anaplasma phagocytophilum* prevalence in small mammals as a function of landscape variables and species abundance variables. The *A. phagocytophilum* prevalence, expressed as the frequency of infected individuals per site, in all small mammals (**a** and **b**), in wood mice (**c** and **d**), and in bank voles (**e**, **f**, **g** and **h**). Fitted binomial regression curves, according to the best multiple explanatory variables binomial Generalized Linear Models, are shown with 95% confidence intervals (light grey, except **h**). Prevalence is shown as a function of bank vole abundance (**a**, **c**, and **e**), as a function of the proportion of wooded habitats in 50 m-radius zones (**b, f** and **h**) or in the trapping habitats (**d**), and as a function of the proportion of ecotones in 200 m-radius zones (**g** and **h**). Disks areas are proportional to samples sizes (see Table 2).

### *Results for* B. burgdorferi *s.l*

The analyses of *B*. *burgdorferi* s.l. prevalence were based on the PCR results for 450 wood mice, 147 bank voles, four field voles, two common pine voles and three crowned shrews. Fifteen wood mice, ten bank voles and one common pine vole were positive (**Table 2**). The numbers of positive wood mice and bank voles per site were not significantly correlated neither was the prevalence (p > 0.05). The results of the selection procedure of the GLMs of *B*. *burgdorferi* s.l. prevalence as a function of landscape variables and species abundance variables are detailed in **Table 4**.

**Table 4:** Best binomial Generalized Linear Models of *Borrelia burgdorferi* s.l. prevalence in small mammals as a function of landscape variables and species abundance variables according to the AICc-based selection procedure.

| All small mammals |  |  |  |  |  |
| --- | --- | --- | --- | --- | --- |
| Model set | Rank | Variables and their significativity | AICc | ΔAICc | R<sup>2</sup> |
| Null model |  |  | 56.7 |  |  |
| Landscape | 1 | **Ecotones.50m\*** | 55.2 | 0.00 | 0.201 |
|  | 2 | Ecotones.100m° | 55.4 | 0.25 | 0.188 |
| Community | None |  |  |  |  |
| Wood mice (*Apodemus sylvaticus*) |  |  |  |  |  |
| Model set | Rank | Variables and their significativity | AICc | ΔAICc | R<sup>2</sup> |
| Null model |  |  | 43.0 |  |  |
| Landscape | None |  |  |  |  |
| Community | None |  |  |  |  |
| Bank voles (*Myodes glareolus*) |  |  |  |  |  |
| Model set | Rank | Variables and their significativity | AICc | ΔAICc | R<sup>2</sup> |
| Null model |  |  | 35.2 |  |  |
| Landscape | 1 | **Ecotones.100m\*** | 33.3 | 0.00 | 0.252 |
|  | 2 | **Ecotones.local\*** | 33.6 | 0.30 | 0.235 |
|  | 3 | **Woodes.local\*** | 33.6 | 0.31 | 0.234 |
|  | 4 | **Wooded.50m\*** | 33.7 | 0.40 | 0.229 |
|  | 5 | dPC.250m° | 34.0 | 0.65 | 0.214 |
|  | 6 | Ecotones.50m° | 34.0 | 0.67 | 0.213 |
|  | 7 | dPC.25m° | 34.0 | 0.68 | 0.212 |
|  | 8 | Wooded.100m° | 34.1 | 0.78 | 0.206 |
|  | 9 | dPC.50m° | 34.1 | 0.79 | 0.206 |
|  | 10 | dPC.100m° | 34.1 | 0.81 | 0.205 |
|  | 11 | dPC.250m + Ecotones.100m | 34.2 | 0.85 | 0.350 |
|  | 12 | dPCflux.250m° | 34.2 | 0.86 | 0.202 |
|  | 13 | dPC.Mg° | 34.3 | 0.94 | 0.197 |
|  | 14 | dPCconnector.250m° | 34.3 | 0.97 | 0.196 |
|  | 15 | Ecotones.100m + Wooded.50m | 34.3 | 1.01 | 0.347 |
|  | 16 | log(Area)° | 34.3 | 1.02 | 0.193 |
| Community | None |  |  |  |  |

Only models with ΔAICc < 2 are shown. Models with connectivity variables that performed worse than models with the wooded patch area have been discarded. Significant codes are: "\*\*\*\*" when $p \leq 0.001$ ; "\*\*\*" when $0.001 < p \leq 0.01$ ; "\*\*" when $0.01 < p \leq 0.05$ ; "°" when $0.05 < p \leq$ ; otherwise, $0.1 < p$ . Significant variables at $p < 0.05$ are in bold.

For all small mammals, the only significant variable in GLMs of *B*. *burgdorferi* s.l. prevalence included the proportion of ecotones in 50 m-radius zones, displaying a significant positive relationship (**Figure 2a**). For prevalence in wood mice separately, no landscape variable displayed a significant relationship. The most supported GLM of prevalence in bank voles included only the proportion of ecotones in 100 m-radius zones, displaying a significant positive relationship (**Figure 2b**). This variable computed in the trapping habitats was also significant. The proportion of wooded habitats in the trapping habitats and in 50 m-radius zones displayed a significant negative relationship (not shown). No species abundances variable was significantly related to any prevalence.

**Figure 2:**
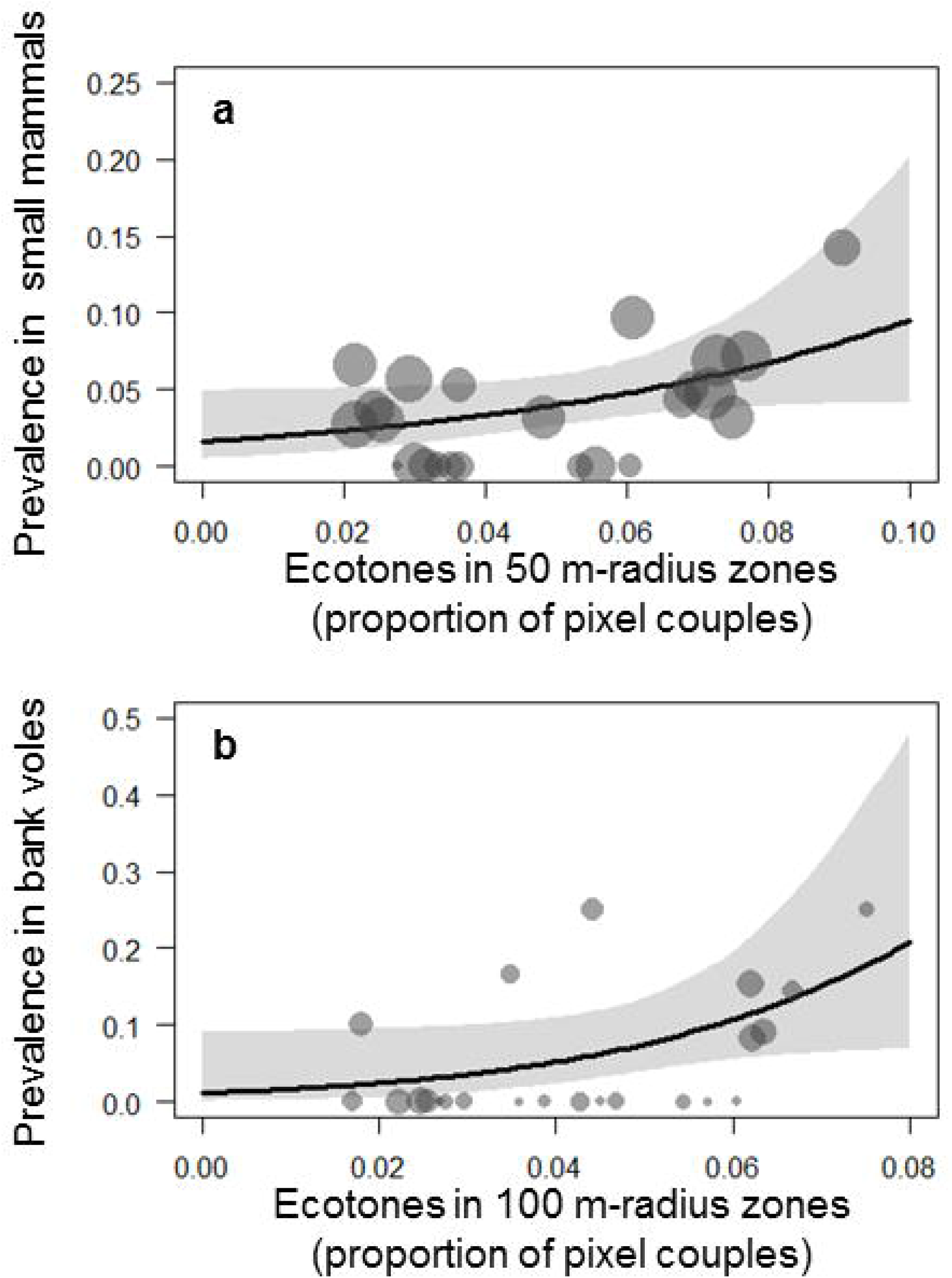
The *Borrelia burgdorferi* sensu lato prevalence in small mammals as a function of landscape variables. The *B. burgdorferi* s.l. prevalence, expressed as the frequency of infected individuals per site, in all small mammals (**a**), and in bank voles (**b**). Fitted binomial regression curves, according to the best single explanatory variable binomial Generalized Linear Models, are shown with 95% confidence intervals (light grey). Prevalence is shown as a function of the proportion of ecotones in 50 m-radius zones (**a**) and in 100 m-radius zones (**b**). Disks areas are proportional to samples sizes (see Table 2).

## Discussion

In this study, we investigated the possible associations between landscape structure, abundances of small mammal species and the prevalence of two tick-borne infectious agents in small mammals, to better understand the spread of these infectious agents in the landscape. The small mammal community at the woodland/hedgerows-grassland ecotones and in forest sites was dominated by the wood mouse and the bank vole, two rodent species with different habitats use, and as such interesting to compare at this landscape scale. However, we found only that the wood mouse abundance was positively correlated to the proportion of ecotones in 500 m-radius zones. This finding is consistent with the propensity of this generalist species to use various complementary habitats (Tew and Macdonald 1994; Ouin et al. 2000). Despite bank vole abundance was not associated to any landscape variable here, in a previous study considering seasonal variation, negative associations with two landscape structural connectivity variables was observed in spring (Perez et al. 2016). Such relationships, also reported in several other studies, are likely caused by a released predation pressure in fragmented landscapes (Szacki 1987; van Apeldoorn et al. 1992; Paillat and Butet 1996; Michel et al. 2007).

### *Prevalence of* Anaplasma phagocytophilum

The numbers of positive individuals as well as the prevalence in wood mice and bank voles were correlated, suggesting a common infection factor and/or an increased exposition of one species in the presence of the other. We observed a higher prevalence in bank voles and a positive, despite weak, relationship between bank vole abundance and the prevalence in all small mammals and in species separately. Although not related to landscape variables, these results support an effect of relative host species abundances on prevalence (**H2**): bank vole likely acts as an amplification host species for the transmission of these bacteria, while wood mouse acts, comparatively, rather as a dilution host species (Rosso et al. 2017; Perez et al. 2017).

Several studies demonstrated that the prevalence of directly transmitted infectious agents increased with host population size, and decreased with population isolation as a consequence of a higher local extinction risk (Begon et al. 2003; Guivier et al. 2011). Here, the proportion of wooded habitats in the surrounding landscape was positively associated to *A*. *phagocytophilum* prevalence in all small mammals and in species separately. More wooded landscapes probably support larger and more connected populations of bank voles resulting in a lower local extinction probability in a given habitat patch (Paillat and Butet 1996). Consequently, a lower local extinction probability of their parasites (including small mammal specialist tick species like *I*. *trianguliceps* and *I*. *acuminatus*), and of the infectious agents they could transmit is expected (**H1**).

### *Prevalence of* Borrelia burgdorferi *sensu lato*

The *B*. *burgdorferi* s.l. prevalence in small mammals, particularly in bank voles, was positively related to the proportion of ecotones within 100 m. These results suggest an enhanced transmission of these bacteria at the interface between wooded and open (crops and grassland) habitats and therefore in fragmented landscapes where such ecotones are frequent. A possible explanation of increased *B*. *burgdorferi* s.l. prevalence in such landscapes is an increased density of small mammals and other hosts, causing an increased density of *I*. *ricinus* nymphs, resulting in an amplification of the transmission cycle (Agoulon et al. 2012; Li et al. 2012; Cayol et al. 2018). However, in previous studies, no relationship was observed between the abundance of *I*. *ricinus* nymphs and *B*. *burgdorferi* s.l. prevalence, and tick density was generally higher in woodlands (Boyard et al. 2008; Hoch et al. 2010; Perez et al. 2017). An alternative explanation is an increased density of hosts competent for the same geno-species than small mammals (Vourc’h et al. 2008).

The mobility of the wood mouse and its ability to use crops and grassland could explain the absence of spatial pattern in the *B*. *burgdorferi* s.l. prevalence of this species. Several studies showed that wood mice, more prone to host *Ixodes* sp. larvae than bank voles, may yield more nymphs infected by those bacteria despite displaying lower bacterial loads (Humair et al. 1993, 1999). Thus, wood mice might act as medium distance dispersers (at least up to 500 m) of infected *Ixodes* sp. larvae that become infective nymphs, as previously suggested in other studies (Boyard et al. 2008; Gassner et al. 2008). These results support the role of host specificity of infectious agents on their distribution patterns (**H3**).

For both infections, prevalence models were not improved by using wooded habitats connectivity measures compared to simpler landscape variables. In our study area, variations in wooded habitats connectivity were maybe not contrasted enough to be captured (Michel et al. 2007). Alternatively, within each considered land cover, the influence of habitat quality and heterogeneity on landscape connectivity for small mammals might have been overlooked and should be accounted for in higher resolution models and/or habitat quality based landscape connectivity models (Mortelliti et al. 2010). It would also be interesting to account for other potential factors affecting small mammal communities and their tick-borne infection prevalence, like predation pressure (Hofmeester et al. 2017).

Finally, species which were rare in the sampled habitats (crowned shrew, field vole and common pine vole) can also host *A*. *phagocytophilum* and/or *B*. *burgdorferi* s.l. (Bown et al. 2008, 2009, 2011). Further studies including habitats where these species can be more abundant (e.g. fallow lands, fields’ margins, clear cuts) and focusing on more hosts species, especially for *B*. *burgdorferi* s.l., would allow a better overall view of the distribution and the spread of these infectious agents in the landscape.

### Conclusion

The *A. phagocytophilum* prevalence could be substantially explained by a positive association with the proportion of wooded habitats in the surrounding landscape at least up to 500 m, validating the host population size hypothesis (**H1**). The *B*. *burgdorferi* s.l. prevalence could be explained only by the presence of ecotones within 100 m. Despite a weak positive relationship between bank vole abundance and *A*. *phagocytophilum* prevalence, no indirect effect of landscape variables on this prevalence has been detected, questioning the overall host community competence hypothesis (**H2**). Indeed, only wood mice abundance was significantly linked to our landscape variables, but at larger scales than the relationships between landscape variables and the prevalence. The prevalence of the two infections displayed contrasted spatial patterns, a difference which likely results from the wider range of hosts and vectors of *B*. *burgdorferi* s.l., as predicted by the infectious agent specificity hypothesis (**H3**). As a whole, our results demonstrate the interest of integrating complementary approaches, such as landscape ecology and community ecology, to better understand the dynamics and the spatial distribution of tick-borne infectious agents to prevent epidemiological risks at the scale of agricultural landscapes.

## Authors’ contributions

GP, SB, AA, GV, OP and AB designed the study. GP, SB, AA, YR, OP and AB participated to the small mammal field sampling. AC performed most of the DNA extractions and the molecular analyses. YR managed the GIS data. GP performed all data analyses and drafted the manuscript. All authors read, commented and approved the manuscript.

## Acknowledgement

We are very grateful to Agnès Bouju, Floriane Boullot, Axelle Durand, Mathieu Gonnet, Olivier Jambon, Maggy Jouglin, Emmanuelle Moreau, Pranav Pandit, and Ionut Pavel who helped in sampling and preparing small mammal tissues before molecular analyses; to Séverine Barry, Amélie Cohadon, Angélique Pion, and Valérie Poux who helped in the lab for the molecular detection of infectious agents; and to Nelly Dorr and Isabelle Lebert who managed the data base of the OSCAR project (https://www6.inra.fr/oscar/). We thank the ‘Zone Atelier Armorique’ (https://osur.univ-rennes1.fr/za-armorique/) for providing the GIS data and for the access to its field facilities. We thank Henri Lemercier and Benoît Chevallier from the ‘Office National des Forêts’ for facilitating the access to the Villecartier forest. The ‘Tiques et Maladies à Tiques’ team of the ‘Réseau Ecologie des Interactions Durables’ group, supported by the INRA and the CNRS gave a rich thinking environment.

This work was funded by the French National Research Agency (ANR-11-Agro-001-04; call for Proposal ‘Agrobiosphere’, OSCAR project). This work is part of the PhD of GP, which was supported by a fellowship from the Brittany region, France. The funders had no role in study design, data collection and interpretation, or the decision to submit the work for publication.

